# Traditionally used aqueous herbal extracts for respiratory ailments exhibit anti-inflammatory activity

**DOI:** 10.1101/2020.12.02.408351

**Authors:** Priya Kannian, Sambasivam Mohana, Manivasagam Vishwarohini, Abdul Rahim Mohamed Saleem

**Affiliations:** Department of Clinical Research, VHS Hospital, Chennai, Tamil Nadu, India; Department of Virology, King Institute of Preventive Medicine and Research, Chennai, Tamil Nadu, India; Department of Food Chemistry & Food Processing, Loyola College, Chennai, Tamil Nadu, India

**Keywords:** Aqueous extracts, acetone extracts, ethanol extracts, immunomodulatory properties, antibacterial activity

## Abstract

**Background:** Phytoconstituents from plants provide a huge array of medicinal properties and have been used for many centuries as household medicines in India to treat respiratory illnesses. There are limited studies on comprehensive evaluations of these plant extracts to understand their healing properties.

**Aim:** To evaluate the antimicrobial and immunomodulatory properties of the 10 commonly used medicinal plants to treat respiratory illnesses in a comprehensive manner.

**Materials & Methods:** The acetone, ethanol and aqueous extracts from the 10 plants were evaluated for antiviral, antifungal and antibacterial activity. Additionally, the aqueous extracts were also evaluated for immunomodulatory properties.

**Results:** None of the extracts exhibited antiviral activity against the four influenza viruses, antifungal activity against *Candida albicans* or antibacterial activity against Gram negative bacilli. Eight acetone extracts, nine ethanol extracts and one aqueous extract *(NSAq)* exhibited antibacterial activity at concentrations below 1mg/ml against *S. pyogenes* and/or *S. pneumoniae.*All the aqueous extracts exhibited dose-dependent anti-inflammatory activity against the innate immunity cytokines, TNF-α and IL-1β, and the Th1 cytokine, IFN-γ, but not against the Th2 cytokine, IL-4.

**Conclusions:** Overall the study elucidates that the acetone and ethanol extracts exhibit antibacterial activity, while the commonly consumed aqueous extracts ameliorate respiratory illnesses through immune modulation.

## 1. Introduction

With the emerging drug resistance among respiratory pathogens like influenza virus, Pneumococci, *Staphylococcus aureus, Klebsiella* spp., etc, and the lack of treatment for some pathogens like the highly pathogenic human corona viruses, new drug discoveries are essential. Plants have always been a good source of chemical compounds with antimicrobial activity. Plants synthesize many types of aromatic substances that are largely secondary metabolites (Cowan, 1999). These substances provide defense against predation from microorganisms, insects and herbivores. Over the years many of them have been adopted as part of human food like chilli peppers, herbs and spices, etc. Major classes of antimicrobial compounds from plants include simple phenols, flavanoids, terpenoids, alkaloids, polypeptides, etc (Cowan, 1999).

In India, traditional household remedies for respiratory and gastrointestinal ailments using locally grown aqueous plant extracts is a common practice. These remedies are based upon the symptoms and not the aetiologies. So, a cough or cold will be treated with aqueous extracts from *Anisochilus carnosus, Adhatoda justacia, Piper betle,* etc irrespective of the underlying bacterial, fungal or viral aetiology (Govindan *et al.* 2004; Khare, 2007; Govindarajan *et al.* 2008; Karunamoorthi *et al.* 2012). Such home remedies are primarily aqueous extracts of the plants. However, most of the phytochemicals being aromatic compounds, organic solvent extractions give a better yield for laboratory testing. Therefore, a pragmatic search for new compounds would be to evaluate the organic solvent extracts as well as the aqueous extracts of the plants commonly used in a comprehensive manner in order to understand their antimicrobial properties.

It is intriguing that these aqueous plant extracts ameliorate disease symptoms even though they have a lower yield of the phytoconstituents, compared to the organic solvent extracts. The likely explanation for this is the potential immunomodulatory properties in these extracts. Thus, additionally, evaluation of the aqueous extracts of the plants for their immunomodulatory properties will greatly help move the field forward in identification of novel drug candidates as well as in elucidation of the healing properties of these medicinal plants.

In the current study, we propose to analyse the antibacterial, antifungal, antiviral and immunomodulatory properties of 10 traditionally used medicinal plants in a comprehensive manner using the acetone, ethanol or aqueous extracts. The findings of this study show that the aqueous extracts exhibit immunomodulatory properties rather than direct antimicrobial activity. While some of the organic solvent extracts exhibited antibacterial activity, thereby providing leads to explore new drug possibilities.

## 2. Materials and Methods

### 2.1 Microbial strains, cell cultures and plant extracts

The four ATCC strains of influenza virus AH_1_N_1_ pdm09 (2009 pandemic strain; A/California/07/2009), AH_3_N_2_ (A/Hong Kong/1/68), BV (Influenza B/Victoria/2/87) and BY (Influenza B/Yamagata/16/88) were obtained from National Institute of Virology (NIV), Pune, India. The ATCC Gram positive cocci - *Streptococcus pyogenes* (ATCC19615) and *Streptococcus pneumoniae* (ATCC49619), Gram negative bacilli - *Escherichia coli*(ATCC10536), *Klebsiella pneumoniae* (ATCC13883) and *Pseudomonas aeruginosa* (ATCC 10145), and *Candida albicans* (ATCC26790) were obtained from HiMedia, India. Human peripheral blood mononuclear cells (PBMCs) were isolated from donor buffy coat bags obtained from VHS Blood Bank, Chennai, India. The Vero cells for the maintenance of influenza virus stocks were obtained from National Centre for Cell Science (NCCS), Pune, India.

The ten plants evaluated in this study were identified taxonomically by a botanist at the herbariums of Southern Regional Centre, Ministry of Environment and Forests, Coimbatore, India, and Plant Anatomy Research Centre, Institute of Herbal Science, Chennai, India. The botanical details of these plants have been described previously (Kannian *et al.* 2020). The names of these 10 plants are listed in table 1. The acetone, ethanol and aqueous extracts were prepared as described previously (Kannian *et al.* 2020). Briefly, the respective plant parts were washed, dried and powdered. The phytoconstituents were extracted and lyophilized. The stock concentrations were prepared using the lyophilized material in Dimethyl sulfoxide (DMSO; HiMedia, India) or hot water. The final yield of the extracts was about 10-20% of the starting material. The phytochemical constituents present in these 10 acetone, ethanol and aqueous extracts were determined by previously described methods (Kannian *et al.* 2020).

### 2.2 Haemagglutination inhibition assays

For the haemagglutination inhibition (HAI) assays, extracts were screened initially at three 10-fold concentrations of 2mg/ml, 0.2mg/ml and 0.02mg/ml in Dulbecco’s Minimum Essential Medium (HiMedia, India) containing 0.25% of DMSO. All extracts were tested along with negative controls, drug controls, vehicle controls and positive controls. All controls were run in duplicates, and all test concentrations were done in hexaplicates. Each extract was tested in three independent experiments.

### 2.3 Broth dilution method

Broth dilution described previously (Kannian *et al* 2020) was employed to determine the antibacterial and antifungal activity of the acetone, ethanol and aqueous extracts from the 10 plants against *S. pyogenes, S. pneumoniae, E. coli, K. pneumoniae, P. aeruginosa* and *C. albicans.* All extracts were tested in duplicates in three independent experiments.

### 2.4 Immunological assays

Immunomodulatory properties of the 10 aqueous extracts were determined using PBMCs isolated from normal healthy blood donors’ buffy coat bags obtained from VHS Blood Bank using Ficol-Paque (Invitrogen, USA). PBMCs were added into 75cm^2^ tissue culture flasks at 10×10^6^ cells/10ml in RPMI (HiMedia, India) containing 10% foetal bovine serum (FBS; HiMedia, India) and incubated at 37°C with 5% CO_2_ for three hours. Monocytes adhered to the bottom while T cells remained suspended. The T cells were collected, washed and counted. The monocytes were scraped off the flasks using cell scrapers, washed and counted. T cells were seeded in 96 well plates at 1×10^6^ cells/ml in the presence or absence of 2μg/ml of phytohaemagglutinin (PHA; Invitrogen, USA). Monocytes were seeded in 96 well plates at 1.5×10^6^ cells/ml in the presence or absence of 10ng/ml of lipopolysaccharide (LPS; Sigma-Aldrich, USA). The aqueous extracts from the plants were added to the stimulated or unstimulated cells at three different concentrations of 50, 100 and 200 μg/ml. Cell controls, LPS/PHA controls and vehicle controls were included for each donor’s cells. The supernatants from the 96 well plates were collected after four hours from the monocytes and after 24 and 48 hours from the T cells. The supernatants were stored at −20°C until further use. Each assay was done in quadruplicates and each extract was tested in two independent experiments using PBMCs from two different donors. TNF-α and IL-1β cytokines were measured in the supernatants collected from monocytes. IFN-γ was measured in the supernatants collected from the T cells after 24 hours. IL-4 was measured in the supernatants collected after 48 hours from the T cells. All these cytokines were measured by ELISA (Diaclone, France) according to manufacturer’s instructions.

## 3. Results and Discussion

### 3.1 NSAq and the organic solvent extracts exhibited antibacterial activity against Gram positive cocci

A number of South Indian plants are used in the form of aqueous concoctions in most households to alleviate respiratory symptoms. These aqueous concoctions are made using different combinations of herbs. The choice of the plants depends on the symptoms rather than the underlying aetiology. Most phytoconstitutents being aromatic substances are generally extracted better in organic solvents (Cowan, 1999). We have previously shown that among the acetone, ethanol and aqueous extracts of these 10 plants, only the acetone extract of *Anisochilus carnosus* exhibited antibacterial activity against *Staphylococcus aureus* (Kannian *et al.* 2020). Therefore, in this study, we hypothesized that the organic solvent extractions are more likely to exhibit direct antimicrobial activity, while the orally consumed aqueous herbal extracts would potentially exhibit activity on the host immune response rather than the microbial agent. To test this hypothesis the acetone, ethanol and aqueous extracts from the 10 commonly used herbs for respiratory illnesses were evaluated for direct antimicrobial activity against the ATCC viral, bacterial and fungal strains of the common respiratory pathogens in India. The phytochemical constituents (alkaloids, flavanoids and phenols) measured in all the extracts are shown in table 1. Terpenoids were present only in the ethanol extracts of *Adhatoda justicia* and *Anisochilus carnosus.* Tannins were present only in the ethanol extract of *Emblica officinalis.* None of the extracts tested exhibited antiviral activity against any of the four commonly prevalent influenza virus strains in India. None of the extracts exhibited antifungal activity against *C. albicans,* or antibacterial activity against the Gram negative bacteria –*E. coli, K. pneumoniae* and *P. aeruginosa.* Among the Gram positive cocci tested, only the aqueous extract of *Nigella sativa (NSAq)* exhibited antibacterial activity against *S. pneumoniae* at a minimum inhibitory concentration (MIC) of 0.02 + 0.007 mg/ml. The acetone and ethanol extracts that exhibited antibacterial activity against *S. pyogenes* and *S. pneumoniae* are listed in tables 2 and 3, respectively. In this study, crude extract concentrations below 1mg/ml were only considered as significant antimicrobial activity as per the recommendations of Rios and Reico (2005). Thus, nine acetone extracts and eight ethanol extracts exhibited antibacterial activity, while only one aqueous extract, *NSAq* exhibited antibacterial activity against the Gram positive cocci.

Cowan (1999) had detailed in his review that any organic solvent extracts of the medicinal plants contained more phytoconstituents compared to their corresponding aqueous extracts. The organic solvent extracts of some of the plants used in this study have shown antibacterial activity similar to that detailed by Khare (2007). Ahmad *et al* (2013) in their review has shown *Nigella sativa* aqueous and ethanolic extracts to have antibacterial activity against Gram positive and Gram negative bacteria. However, the main difference in our study is due to the cut-off significance fixed at 1mg/ml of the crude extracts for antimicrobial activity. Thus, *NSAq,* the only aqueous extract with direct antibacterial activity could serve as a potential alternative to treat antibiotic-resistant *S. pneumoniae.* Additionally, our findings open venues to explore the organic solvent extracts of the plants that exhibited antibacterial activity for potential phytochemicals that could serve as new drug candidates.

### 3.2 The aqueous extracts exhibited robust anti-inflammatory activity

Since the aqueous extracts hardly exhibited any direct antimicrobial activity, the alternate scientific explanation for a symptomatic relief is immunomodulatory properties. So next, we evaluated the immunomodulatory effect of the 10 aqueous extracts against two innate immunity cytokines - TNF-α and IL-1β, and two adaptive immunity cytokoines - IFN-γ (Th1 cytokine) and IL-4 (Th2 cytokine). The immunomodulatory effect was evaluated using normal human PBMCs. Even though primary cells are highly variable, appropriate cell controls and drug controls will facilitate the determination of the immunological effect. Additionally, since the primary human cells would depict the *in vivo* activity, these primary cells are more reliable for such experiments.

After four hours of LPS stimulation, the monocytes produced high levels of TNF-α (fig. 1) and IL-1β (fig. 2). In the presence of each of the 10 aqueous extracts, the levels of both these innate immunity cytokines were decreased in a dose dependent manner. A similar trend was shown by the monocytes isolated from PBMCs of two different donors tested in two independent experiments. Additionally, the levels of IFN-γ, a Th1 cytokine decreased in a dose dependent manner in PHA-stimulated T cells (fig. 3). However, there was no effect on the levels of IL-4, a Th2 cytokine secreted by the PHA-stimulated T cells (data not shown). The baseline cytokine levels did not increase in these unstimulated cells in the presence of the aqueous extracts. This suggests that they do not play a role in pro-inflammatory activity. Dose dependent decrease in cytokines indicates that the aqueous extracts from the 10 plants in this study exhibited an antiinflammatory activity against the innate immunity cytokines and Th1 cytokine, but not against the Th2 cytokine.

**Fig. 1:**
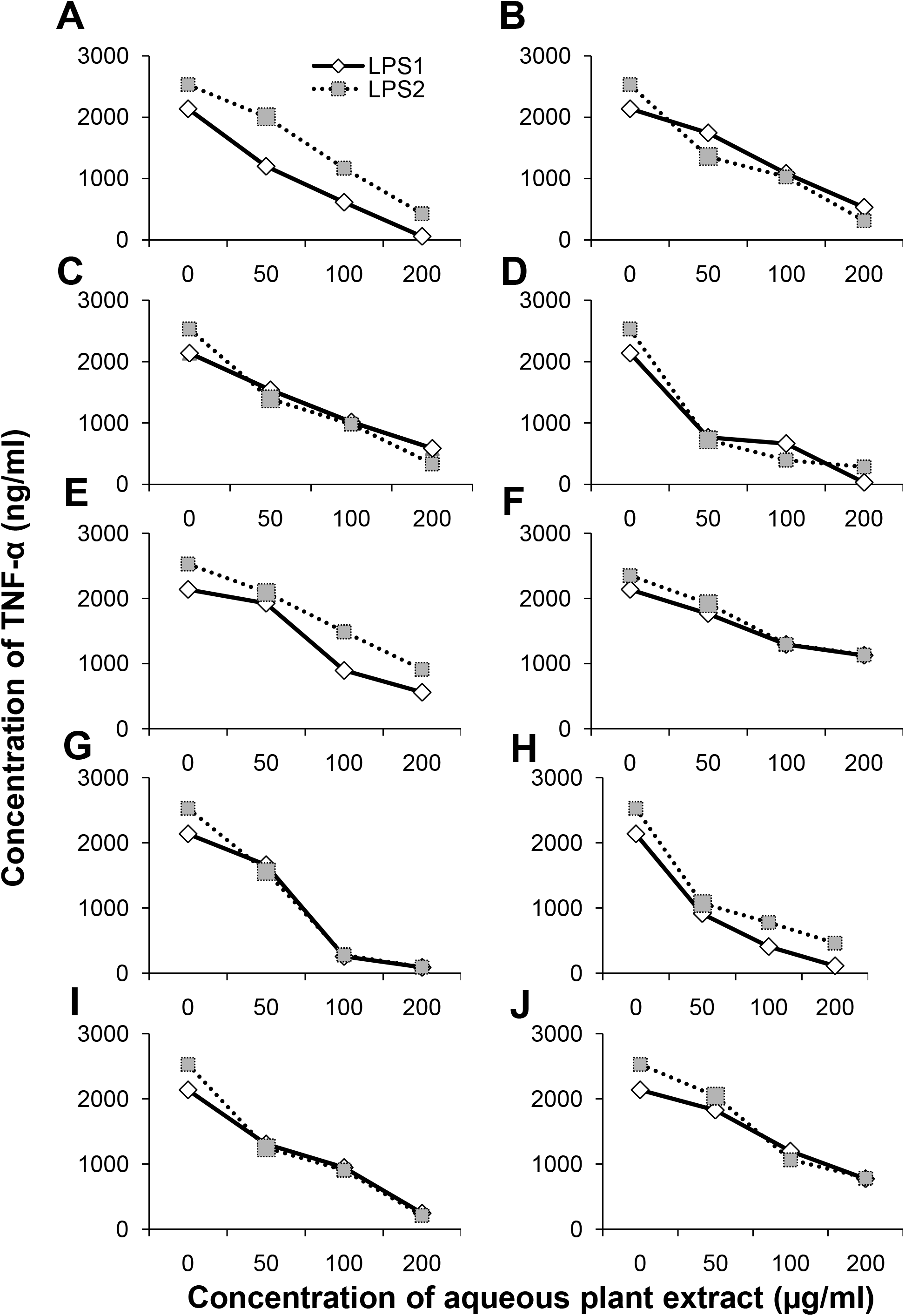
Dose-dependent anti-inflammatory activity of the aqueous extracts from the 10 plants against tumour necrosis factor – alpha (TNF-α). The X-axis denotes the increasing concentrations of the aqueous extracts in μg/ml. The Y-axis denotes the concentrations of TNF-α in pg/ml. The white diamond and solid line represent the lipopolysaccharide (LPS)-induced monocytes of donor 1 (LPS-1). The grey square and dotted line represent the LPS-induced monocytes of donor 2 (LPS-2). A - *Acalypha indica;* B - *Adhatoda justicia;* C - *Anisochilus carnosus;* D - *Emblica officinalis;* E - *Mukia scabrella;* F - *Nigella sativa;* G - *Ocimum sanctum;* H - *Piper betle* black; I - *Piper betle* white; J - *Solanum trilobatum.*

**Fig. 2:**
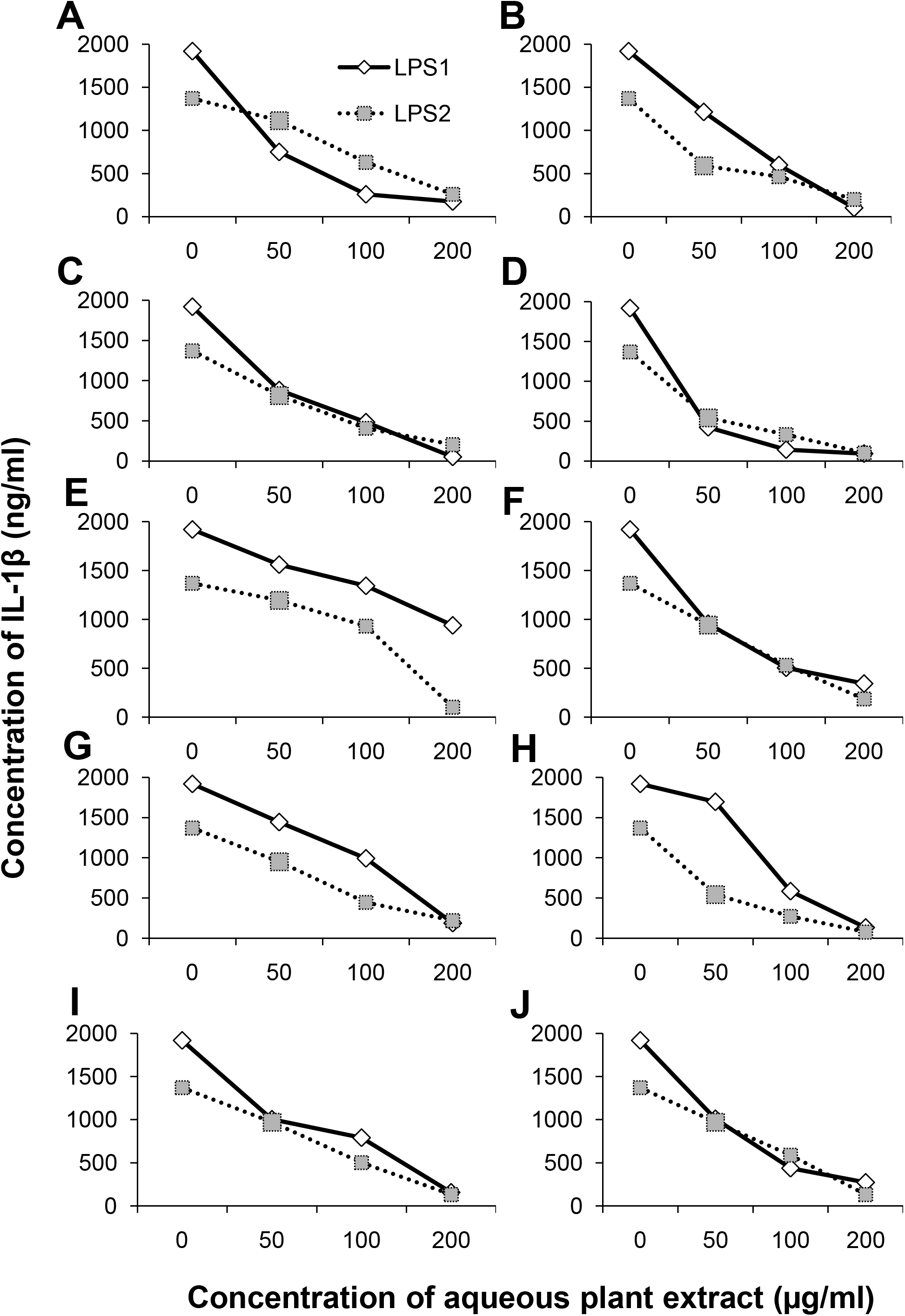
Dose-dependent anti-inflammatory activity of the aqueous extracts from the 10 plants against Interleukin – 1 beta (IL-1β). The X-axis denotes the increasing concentrations of the aqueous extracts in μg/ml. The Y-axis denotes the concentrations of IL-1β in pg/ml. The white diamond and solid line represent the lipopolysaccharide (LPS)-induced monocytes of donor 1 (LPS-1). The grey square and dotted line represent the LPS-induced monocytes of donor 2 (LPS-2). A - *Acalypha indica;* B - *Adhatoda justicia;* C - *Anisochilus carnosus;* D - *Emblica officinalis;* E - *Mukia scabrella;* F - *Nigella sativa;* G - *Ocimum sanctum;* H - *Piper betle* black; I - *Piper betle* white; J - *Solanum trilobatum.*

**Fig. 3:**
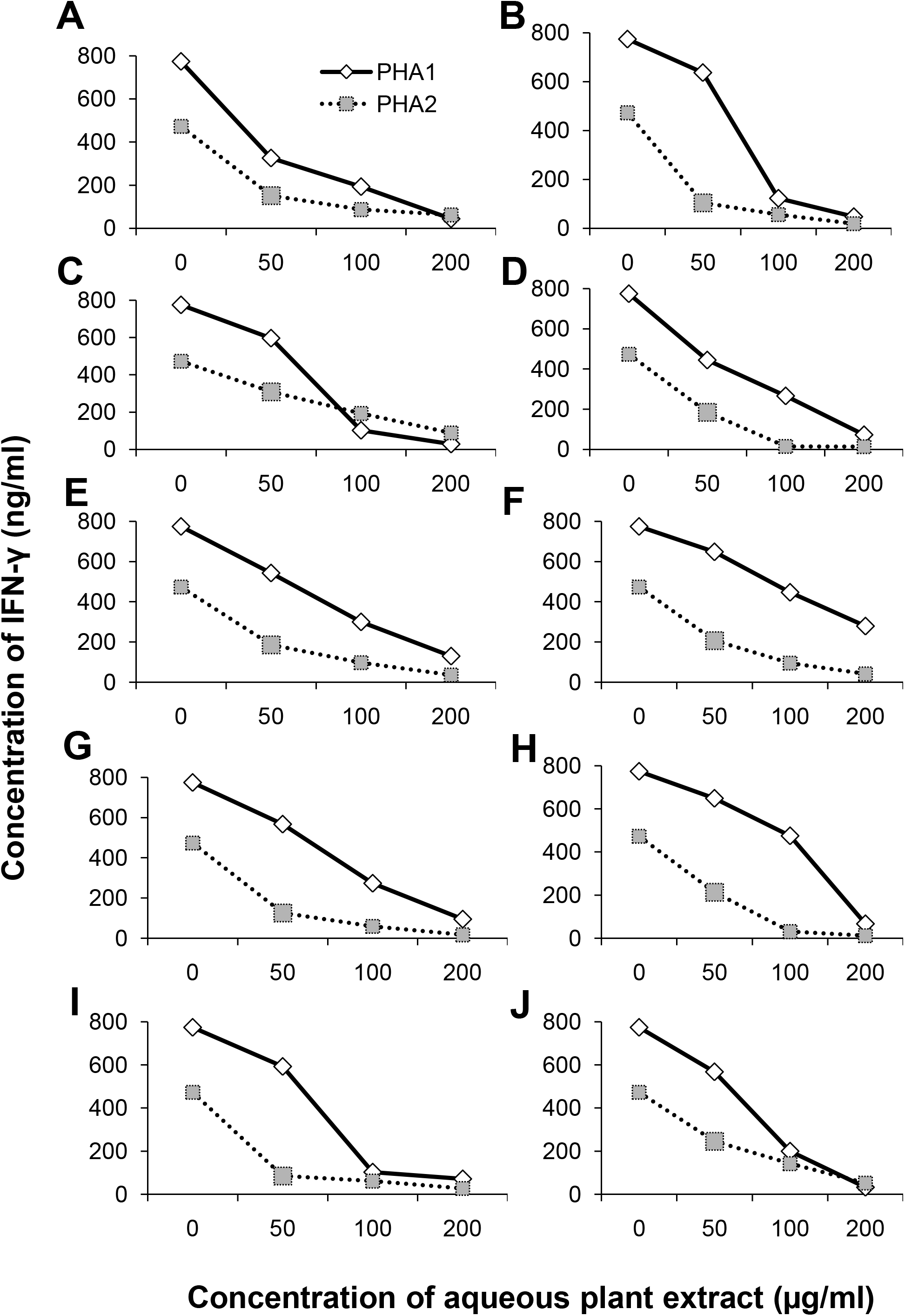
Dose-dependent anti-inflammatory activity of the aqueous extracts from the 10 plants against Interferon – gamma (IFN-γ). The X-axis denotes the increasing concentrations of the aqueous extracts in μg/ml. The Y-axis denotes the concentrations of IFN-γ in pg/ml. The white diamond and solid line represent the phytohaemagglutinin (PHA)-induced T cells of donor 1 (PHA-1). The grey square and dotted line represent the PHA-induced T cells of donor 2 (PHA-2). A - *Acalypha indica;* B - *Adhatoda justicia;* C - *Anisochilus carnosus;* D - *Emblica officinalis;* E - *Mukia scabrella;* F - *Nigella sativa;* G - *Ocimum sanctum;* H - *Piper betle* black; I - *Piper betle* white; J - *Solanum trilobatum.*

Kim *et al* (2020) and Wu *et al* (2018) have previously shown the anti-inflammatory activity of the methanol extract of *Acalypha australis* and the polyphenol enriched fraction of *Acalypha wilkesiana,* respectively in mice. Our study revealed a similar anti-inflammatory activity in the aqueous extracts of the Indian variety of the same plant. Similarly, anti-inflammatory activity of a variant plant species or using a different solvent extract has been shown for few of the plants tested in our study - *Adhatoda justicia* (Nunes *et al* 2018), *Ocimum sanctum* (Choudhury *et al* 2014) and *Piper betle* (Pandey *et al* 2010). The aqueous extract of *Nigella sativa* has been shown to decrease the production of pro-inflammatory cytokines like TNF-α and IL-6 using splenocytes of BALB/c and C57/BL6 mice.

## 4. Conclusions

Overall, this study clearly indicates that the aqueous extracts of the 10 plants used in this study exhibit an anti-inflammatory activity on the innate immunity and Th1 cytokines. These aqueous extracts did not show any direct antimicrobial effect against the common respiratory pathogens, except *NSAq.* In South India, combinations of these aqueous concoctions are commonly used as household remedies for respiratory ailments, irrespective of the underlying aetiologies. Many of the acetone and ethanol extracts have shown antibacterial activity. Further pharmacological investigations to isolate and identify these new phytoconstituents that can act upon drug resistant bacteria are warranted. This study comprehensively elucidates that these aqueous extracts ameliorate the respiratory symptoms through immune modulation rather than direct antimicrobial effect.

## Author contributions

PK was involved in conceiving, designing, execution and interpretation of the study as well as writing of the manuscript. SM, MVR and ARMS were involved in the execution of different parts of the study.

## Abbreviations

ATCC: American Type Culture Collection
pdm: pandemic
PBMCs: peripheral blood mononuclear cells
NCCS: National Centre for Cell Science
DMSO: dimethyl sulfoxide
HAI: haemagglutination inhibition
CO_2_: carbon dioxide
RPMI: Roswell Park Memorial Institute
FBS: foetal bovine serum
PHA: phytohaemagglutinin
LPS: lipopolysaccharide
TNF-α: tumor necrosis factor-alpha
IL-1β: interleukin 1-beta
IFN-γ: interferon-gamma
IL-4: interleukin 4
ELISA: enzyme linked immunosorbent assay
NSAq: *Nigella sativa* aqueous

